# Towards the Genome-scale Discovery of Bivariate Monotonic Classifiers

**DOI:** 10.1101/2023.02.22.529510

**Authors:** Océane Fourquet, Martin S. Krejca, Carola Doerr, Benno Schwikowski

## Abstract

**Motivation:** Bivariate monotonic classifiers (BMCs) are based on pairs of input features. Like many other models used for machine learning, they can capture non-linear patterns in high-dimensional data. At the same time, they are simple and easy to interpret. Until now, the use of BMCs on a genome scale was hampered by the high computational complexity of the search for pairs of features with a high leave-one-out performance estimate.

**Results:** We introduce the *fastBMC* algorithm, which drastically speeds up the identification of BMCs. The algorithm is based on a mathematical bound for the BMC performance estimate while maintaining optimality. We show empirically that fastBMC speeds up the computation by a factor of at least 15 already for a small number of features, compared to the traditional approach. For two of the three clinical datasets that we consider here, the resulting possibility of considering much larger sets of features translates into significantly improved classification performance. As an example for the high degree of interpretability of BMCs, we discuss a straightforward interpretation of a BMC glioblastoma survival predictor, an immediate novel biomedical hypothesis, options for biomedical validation, and treatment implications.

**Conclusions:** fastBMC enables the rapid construction of robust and interpretable ensemble models using BMC, facilitating the discovery of interesting gene pairs and their contributions to the underlying biology.

**Availability:** We provide the first open-source implementation for learning BMCs, and an implementation of fastBMC in particular, all in Python, at https://github.com/oceanefrqt/fastBMC.

## 1 Introduction

High-throughput RNA sequencing and new data analysis methods show great potential for the diagnosis and classification of disease [1, 2]. In particular, supervised machine learning methods are powerful tools for discovering associations between molecular profiles and disease states and variants. However, many advanced methods that can model nonlinear associations represent “black box” models that represent complex high-dimensional associations that are challenging to interpret in the light of other biomedical knowledge. Without this interpretability, the generation of testable hypotheses and insight into complex biology is often impossible.

Two complementary approaches can be used to improve interpretability in this context. Firstly, feature selection and dimensionality reduction methods can be used to identify the most relevant features and filter out noise. However, it is essential to note that most common feature selection methods, such as filtering by variance or correlation, analyze features independently and do not take into account the potentially more complex associations that can be identified by downstream machine learning models. Secondly, using simpler data patterns in models can facilitate easier understanding and translation into biological or clinical contexts.

One class of machine learning methods with intrinsically better interpretability are bivariate classifiers, which involve only pairs of features. These classifiers are more complex than univariate classifiers but are still simple enough to allow effective learning on small datasets. They also allow for graphical representation and analysis of the models and data in two dimensions. For example, bivariate tree methods [3] are able to determine decision trees based on pairs of features with good predictive power. Top-scoring pair classifiers [4, 5], are based on pairs of features whose relative magnitude is associated with the prediction target. Another class of models that offer a unique balance between complexity and interpretability are bivariate monotonic classifiers [6]. They are more complex than Top-scoring pair classifiers, as they consider more than a simple linear separation function but are more constrained than bivariate tree methods, as they enforce a monotonic relationship between the features and the class label. This monotonicity pattern enables the models to be easily understood and visualized, making them highly usable for domain experts. We focus on these bivariate monotonic classifiers (BMCs), which have already been successfully used to predict disease severity from an ensemble BMC in the context of dengue fever [6]. However, this previously developed approach, which we call naïveBMC, has limitations, such as computation time, which restricts the size of the input data.

### Contribution

This paper addresses the construction of ensemble BMCs for larger genome-scale feature sets. We propose two key contributions. Firstly, we introduce a *Preselection Algorithm* that accelerates the identification of suitable BMCs (Algorithm 1), based on a lower bound of the BMCs estimate (Theorem 1), producing a new version of naïveBMC called *fastBMC*. Secondly, we provide a reassessment of the efficacy and contributions of BMCs and ensemble BMC for biomedical research.

### Outline

Section 2.1 provides high-level definitions for the classification problem and the ensemble BMC approach (omitted in [6]). In Section 2.2, we introduce our fastBMC approach. We present the results of our empirical evaluation in Section 3, which addresses the question of whether and how the testing of many more feature pairs results in better classification performance, as well as the interpretability. Finally, we present a formal description of the naïveBMC, fastBMC, and Preselection Algorithm while proving its correctness (Section 4.2).

## 2 Methodology

This section provides introductory high-level descriptions of the classification approach introduced by Nikolayeva et al. [6] (Section 2.1), and the idea behind fastBMC (Section 2.2). For a formal description of the methods, refer to Section 4.

### 2.1 Bivariate Monotonic Classifiers and the naïveBMC Approach

Bivariate monotonic classifiers (BMCs) use general monotonic functions as decision boundaries [7]. BMCs are operating on continuous data to predict binary outcomes, offering a straightforward and effective approach to identify relationships between two variables. In particular, as seen in [6] where they were applied to transcriptomic data to predict the severity of dengue infections in Cambodian patients, these classifiers provide easily interpretable results. Hovewer, BMCs work under the assumption of monotonic relationships, which limits the type of discovery.

Figure 1 shows one of the best BMCs for the prediction of dengue severity in the transcriptomic dataset from [6]. Interestingly, the known biological functions of both transcripts overlap in the highly specific context of T cell co-stimulation [8, 9], which is known to be crucial in the response to the dengue virus [10]. Therefore, this classifier can be interpreted as a complex biological association, in which *simultaneously low* levels of OX40 and CD40 ligand transcripts are associated with severe dengue infection.

**Figure 1:**
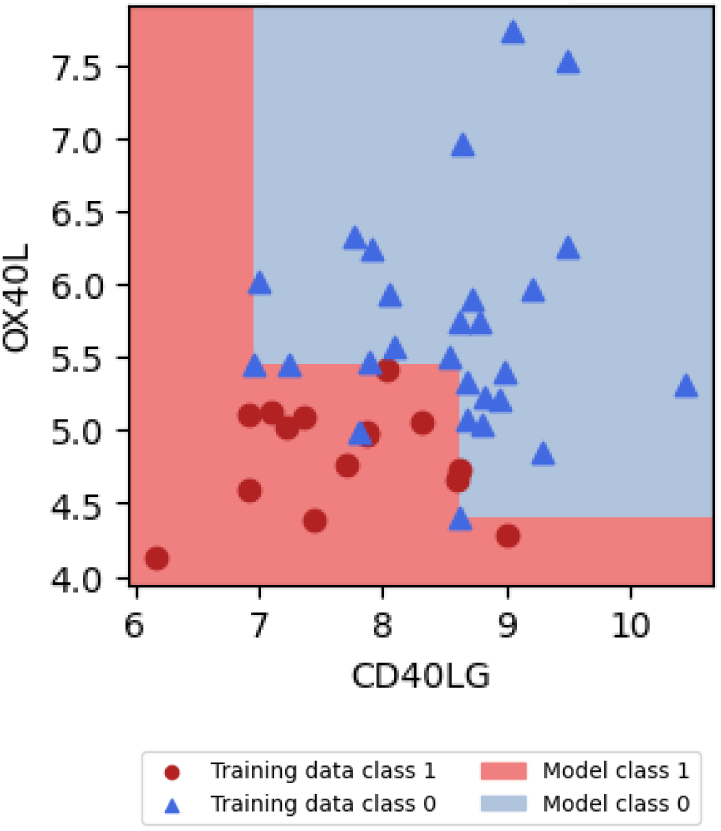
Visualization of a bivariate monotonic classifier. A bivariate monotonic classifier (BMC) is a function *f* : ℝ^2^ → {0, 1} that is monotonic in both dimensions. The function *f* thus divides ℝ^2^ into two classes. This specific BMC, based on OX40 and CD40 ligand transcripts, is an example from [6].

To capture more associations that involve more than two transcripts, multiple BMCs can be combined into an ensemble classifier (ensemble BMC), while constrained not to contain BMCs using the same transcripts (only disjoint BMCs). Nikolayeva et al. [6] applied such an ensemble BMC to the classification of clinical dengue infections into *mild* and *severe*, according to the blood transcriptome at hospital admission. Their approach empirically performed as good as or better than other classifiers.

The ensemble BMC by Nikolayeva et al. [6] was built from BMCs with the best empirical performance estimate, which requires the computation of this estimate for each possible pair of features. As this number grows quadratically with the number of features, this construction (which we refer to as *naïveBMC* ) is computationally expensive and, in practice, can only be used on small subsets of typical genome-scale datasets. Nikolayeva et al. [6] were using two performance estimates for each BMC. The first one corresponds to the error rate when the BMC is trained on all the data (ER_full_). The second one, more robust but also more time-consuming, is the error rate computed with a leave-one-out cross-validation (ER_loocv_). ER_loocv_ is an integral part of the method for building the ensemble BMC.

### 2.2 Preselection and fastBMC Algorithms

As mentioned in Section 2.1, what makes the naïveBMC particularly time-consuming computationally speaking is the calculation of ER_loocv_ for all the BMCs.

We develop a BMC preselection algorithm that identifies the top-performing BMCs according to ER_loocv_, but based primarily on ER_full_, which is much faster to calculate and much less resource-intensive. This algorithm is based on the property that for a given BMC, its ER_full_ is a lower bound of its ER_loocv_ (proof available in Section 4.3). The Preselection Algorithm proceeds in three phases and takes as parameter a value *k*, which corresponds to the number of disjoint BMCs expected to build the ensemble BMC. Phase 1 consists in calculating the ER_full_ for the set of BMCs and ordering them according to their ER_full_ (ascending order). In Phase 2, the ER_loocv_ of the BMCs with the lowest ER_full_ are calculated, until the *k* disjoint BMCs are reached. In this subset of BMCs, the maximum ER_loocv_ observed is used to determine the initial threshold, from which BMCs with a higher ER_full_ are directly eliminated. Phase 3 can then begin. It consists of updating the threshold, continuing the ER_loocv_ calculations (in increasing ER_full_ order) and always selecting the minimal subset containing *k* disjoint BMCs. It stops when the top-performing BMCs are identified. Figure 2 is a graphic illustration of the Preselection Algorithm. A formal description and a pseudocode of the algorithm are available in Section 4.3.1.

**Figure 2:**
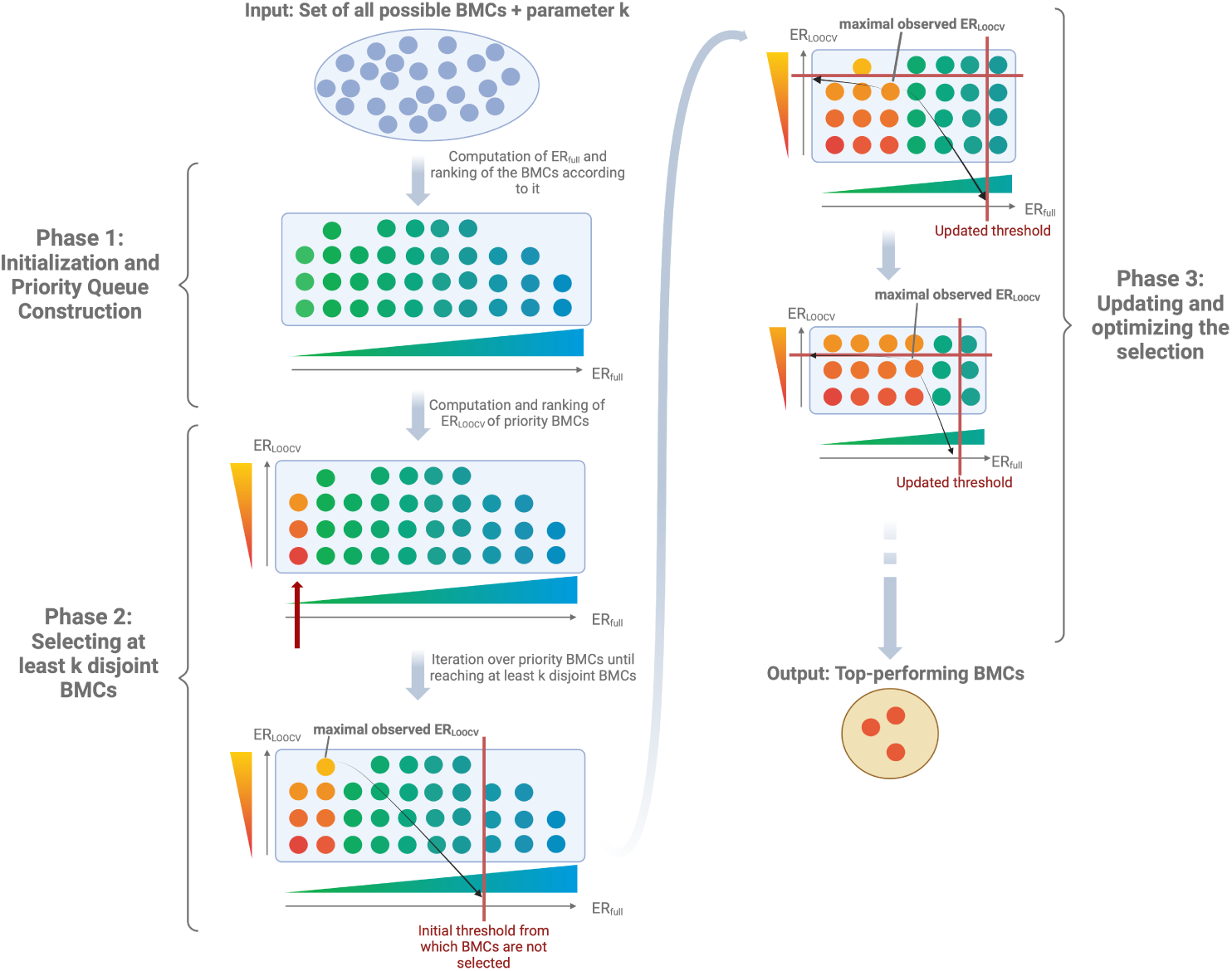
Operation of the Preselection Algorithm for BMCs: Each circle represents a BMC, with the blue-green gradient indicating ER_full_ and the yellow-red gradient representing ER_loocv_. The interplay of colors reveals the computational dynamics within the algorithm. Specifically, the blue-green gradient indicates that all BMCs are initially ranked according to ER_full_. Subsequently, as illustrated by the yellow-red gradient, the ER_loocv_ values are iteratively computed to define and update the threshold.

The fastBMC is an extension of the naïveBMC. It introduces the Preselection Algorithm in the various steps used to build the ensemble BMC.

## 3 Empirical Evaluation

This section provides an overview of the datasets and the experimental settings employed (Section 3.1). The empirical evaluation on clinical datasets that follows assesses the efficiency of fastBMC, focusing on key aspects such as the improvement in running time (Section 3.2), a comparative analysis of its performance relative to both the naïveBMC and other classical algorithms (Section 3.3). Finally, in Section 3.4, we examine of the interpretability of the results on one of the clinical datasets in the light of existing knowledge, hypotheses and outline possible validation options and implications for treatment.

### 3.1 Datasets and Experimental Setup

This section describes the three datasets used in the empirical evaluation, as well as the parameterization of fastBMC.

#### 3.1.1 Description of the Datasets

##### Dengue Dataset

This dataset comes from the study by Nikolayeva et al. [6], in which an RNA signature predicted the severity of dengue in young patients. The set contains data from 42 patients, of whom 15 developed a severe form of dengue and 27 a non-severe form. In Nikolayeva et al. [6], the 2 653 features with the highest variance were used.

##### Leukemia Dataset

This dataset comes from the study by Golub et al. [11], showing how leukemia cases can be classified into acute myeloid leukemia and acute lymphoblastic leukemia. 38 samples were used for training and 34 for testing. The entire data comprises 7 129 features for 72 samples. It is accessible at https://www.kaggle.com/datasets/crawford/gene-expression.

##### Glioblastoma Dataset

This dataset comes from the study by Reifenberger et al. [12] aimed at identifying transcriptomic markers of long-term glioblastoma survival. It comprises 23 samples from long-term (overall survival more than or equal to 36 months) and 16 from short-term survivors (overall survival less than or equal to 12 months). Data were retrieved from the Gene Expression Omnibus (GEO) database at http://www.ncbi.nlm.nih.gov/geo/ (accession number *GSE* 53733). We used only the 16 065 features corresponding to coding genes.

To further reduce the number of features while retaining the most relevant information, we first eliminated those features with an empirical variance below a fixed threshold. Varying this threshold allowed us to [i] generate subdatasets of different sizes, [ii] to profile the speed-up obtained by fastBMC on these subdatasets (Section 3.2), and [iii] to examine whether ability to process larger datasets using fastBMC results in improved classification performance (Section 3.3). In Section 3.4 the variance threshold is fixed so that the resulting subdatasets contain around 1000 transcripts.

#### 3.1.2 Parametrization

The Preselection Algorithm takes as a parameter the number of disjoint BMCs desired for the ensemble model. In the naïveBMC and also in the fastBMC, this parameter is determined in order to maximize the performance of the ensemble BMCbetween a range of 1 and *k_max_* ∈ ℕ. Both naïveBMC and fastBMC take k_max_ as a parameter, i.e., the maximal number of disjoint BMCs that we want to test. In this study, as an example for concrete use case, such as identifying a gene pair signature, a reasonable and practical parameter is *k*_max_ = 20. This constraint is imposed to ensure that the resulting signature remains interpretable and manageable, allowing for meaningful analysis and downstream applications, while also mitigating the risk of overfitting or information overload.

### 3.2 Obtained Speed-up

Figure 3 shows, for each of the datasets, the running times of fastBMC and naïveBMC on the subdatasets of varying size.

**Figure 3:**
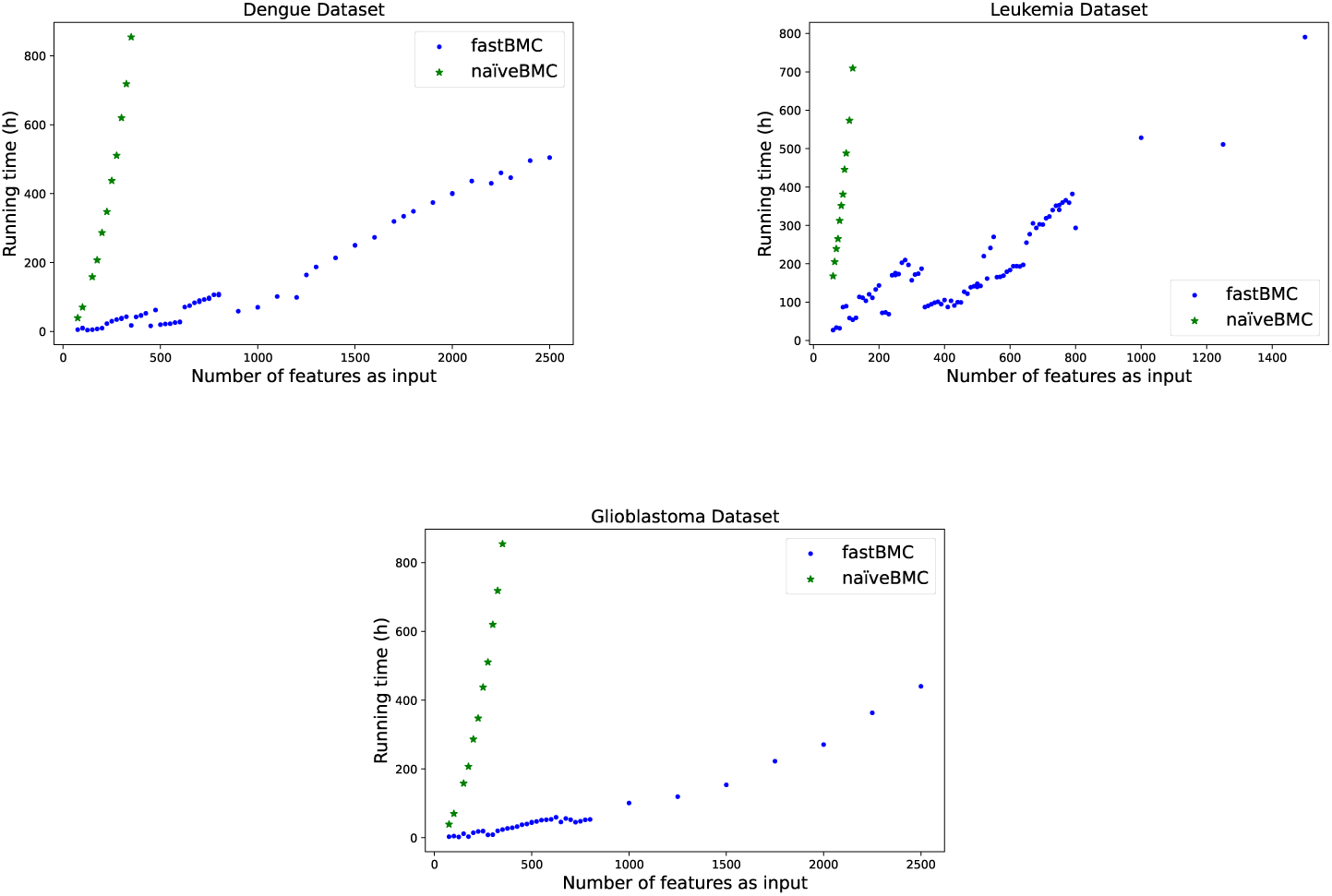
Running times in hours of naïveBMC and fastBMC on subdatasets of varying size, across three clinical datasets.

Note that, interestingly, the running time of the fastBMC does not strictly increase with the number of input features: A newly added feature may lead to a reduction of the threshold *t*, leading fastBMC to perform fewer ER_loocv_ evaluations than without the newly added feature.

More importantly, on subdatasets of 250 or more features, fastBMC is faster by at least a factor of 15 (Table 1), and Figure 3 suggests that this factor increases with increasing subdataset size.

**Table 1:** Comparison of *user running time (URT)* between naïveBMC and fastBMC, on clinical datasets with 250 transcriptomic features. naïveBMC on the leukemia dataset was aborted after 1104 core-hours

| Dataset | URT<br>fastBMC (h) | URT<br>naïveBMC (h) | Speed-up |
| --- | --- | --- | --- |
| Dengue | 30 | 449 | 15x |
| Glioblastoma | 19 | 325 | 17x |
| Leukemia | 170 | — | — |

### 3.3 Does the Speed-up Translate into Better Classifiers?

Firstly, we aim to investigate whether exploring a larger feature space is worthwhile, and if it enables the discovery of more accurate classifiers. Following this initial assessment, we conduct a comparative analysis of the performance of our proposed method, fastBMC, against other classical techniques.

#### 3.3.1 Comparison with naïveBMC

We now turn to the question of whether the increase in the number of features that can be processed by fastBMC allows the discovery of better classifiers. To do this, we compute the AUC score of fastBMC and naïveBMC for varying subdatasets of increasing size across our three clinical datasets. As before, we aborted any analysis that took more than 1104 hours (user running time).

Figure 4 shows the result. First, we see that the performance of naïveBMC and fastBMC for the same input size is generally comparable. The elimination of certain low-performing models by the Preselection Algorithm does not appear to have a strong influence on classification performance. Second, we see that, for both approaches, a larger input size does not necessarily result in better performance. This observation is in line with the overfitting that can also be observed in other machine learning methods when too many features are provided. Third, for the dengue and leukemia datasets, the best performance is achieved for input sizes larger than what naïveBMC can handle within our time limit. However, for the glioblastoma dataset, the best performance is achieved using only a few features, which is within the capabilities of naïveBMC (albeit, using much more computation time).

**Figure 4:**
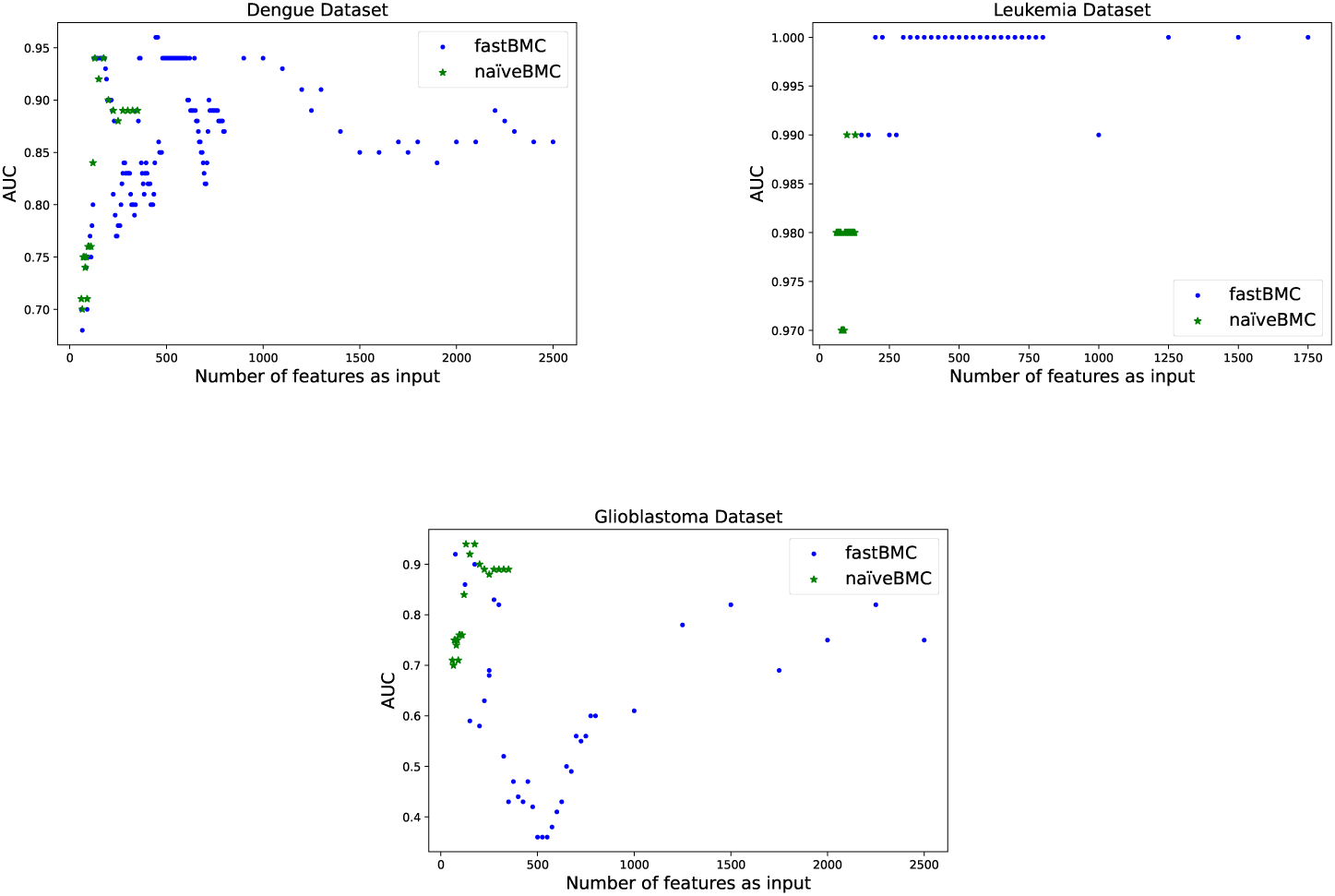
AUC performance on three clinical datasets, of ensemble BMCs constructed using fastBMC or naïveBMC, and for varying numbers of features. curve (AUC), accuracy (Acc), and F1-score (F1). The AUC metric provides an indication of the model’s ability to distinguish between positive and negative classes, while accuracy and F1-score offer insights into the model’s precision and recall. We applied fastBMC to the glioblastoma and leukemia datasets and we opted not to use the dengue dataset, as naïveBMC has already been compared to other methods [6]. To focus on the most informative features while keeping a significant amount of transcripts, we filter both datasets by variance. This results in a subset of 1768 genes for the glioblastoma dataset and 1379 genes for the leukemia dataset.

#### 3.3.2 Comparison with Classical Methods

To evaluate the performance of fastBMC, we compare it to a range of classical classification methods provided by scikit-learn [13]. Specifically, we consider logistic regressions (LR), support vector machines (SVM) with both linear and radial basis function (RBF) kernels (SVM*_linear_* and *SV M_rbf_* ), decision trees (DT), and random forests (RF), all using their default settings. This comparison allows us to assess the effectiveness of fastBMC relative to well-established methods. We use a set of commonly used metrics, including the area under the receiver operating characteristic

For the Leukemia dataset, using fastBMC, we obtained an ensemble BMC containing 9 BMCs, while for the glioblastoma dataset, we obtained an ensemble BMC of 20 BMCs. We evaluated the performance of these ensemble BMCs using a leave-one-out cross-validation (as explained in more details in Section 4.1.3). We performed the same step to evaluate the models constructed with LR, DT, RF, SVM_linear_ and SVM_rbf_, on the thousand genes. We have observed that fastBMC, without being particularly superior to the others, is always comparable to the bests (Table 2).

**Table 2:** Performance of different models on leukemia and glioblastoma datasets. Acc: Accuracy; F1: F1-score; AUC: area under the receiver operating characteristic curve.

| | | $fastBMC$ | $LR$ | $DT$ | $RF$ | $SVM_{linear}$ | $SVM_{rbf}$ |
| --- | --- | --- | --- | --- | --- | --- | --- |
| Leukemia | Acc | 0.97 | 0.96 | 0.81 | 0.97 | 0.97 | 0.94 |
|  | F1 | 0.96 | 0.94 | 0.74 | 0.96 | 0.96 | 0.91 |
|  | AUC | 0.98 | 1 | 0.80 | 1 | 0.99 | 0.99 |
| Glioblastoma | Acc | 0.64 | 0.64 | 0.49 | 0.56 | 0.64 | 0.49 |
|  | F1 | 0.59 | 0.53 | 0.33 | 0.41 | 0.53 | 0 |
|  | AUC | 0.64 | 0.63 | 0.46 | 0.57 | 0.56 | 0.16 |

### 3.4 Interpretability, Hypothesis Generation, Validation Options, and Treatment Implications

BMCs allow prediction, but the interpretation of BMCs in light of existing knowledge may also lead to new insights. Figure 5 shows the glioblastoma BMCs obtained in Section 3.3.2. Some of the genes identified using have already been identified in a previous study using the same dataset [14]. As we illustrate in this section, BMCs may not only be highly predictive, but also represent novel hypotheses that can bring potentially disparate prior knowledge about the two genes into the common functional context of the output variable. The conceptual simplicity of the BMC renders the novel hypotheses highly interpretable and experimentally testable.

**Figure 5:**
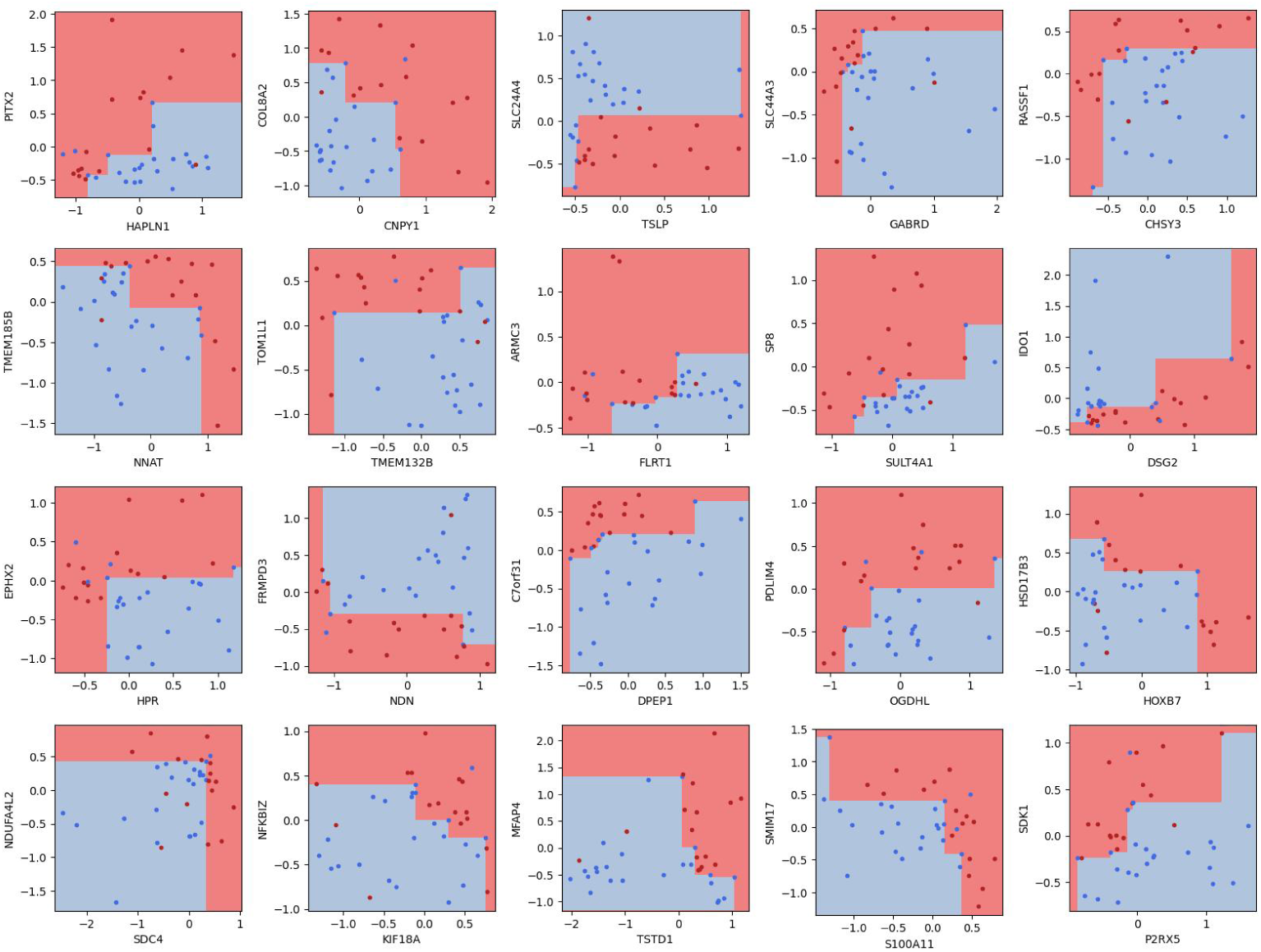
ensemble BMC obtained on the glioblastoma dataset using fastBMC. The dots correspond to the training data and the background colors correspond to the model. The red color is associated with short-term survival and the blue color with long-term survival.

We will discuss in more detail the SDC4/NDUFA4L2 BMC, shown enlarged in Figure 6, which represents an association of simultaneous low expression of both genes with long-term survival.

**Figure 6:**
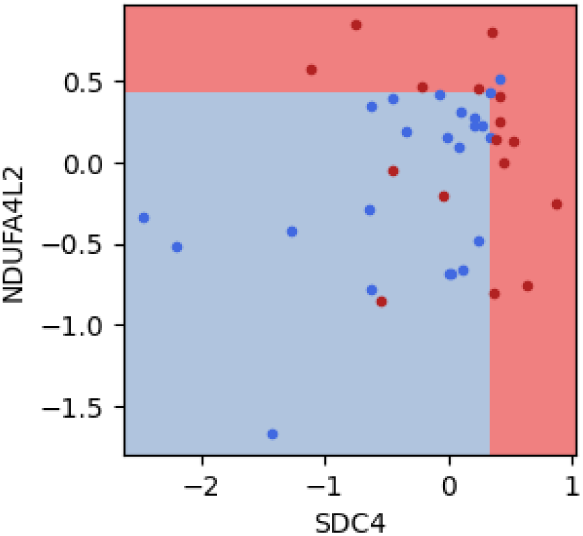
The SDC4/NDUFA4L2 BMC reveals the association of simultaneous low expression of both genes in glioblastomas with long survival.

#### Prior Knowledge in the Glioblastoma Context

SDC4 (Syndecan-4) is a transmembrane heparan sulfate proteoglycan involved in cell–matrix interactions, focal adhesion formation, and the regulation of cell migration and invasion [15]. In glioblastoma, high expression of SDC4 has been associated with enhanced invasive behavior and increased cell motility, which are key drivers of tumor progression [16]. Elevated SDC4 can amplify intracellular signaling cascades, such as those mediated by protein kinase C (PKC) and RhoA, that promote cytoskeletal rearrangements and invasive phenotypes [17, 18].

NDUFA4L2 is a hypoxia-inducible gene encoding a mitochondrial protein associated with complex I of the electron transport chain [19]. Its expression is upregulated under low oxygen conditions, a hallmark of the glioblastoma microenvironment [20]. High levels of NDUFA4L2 are thought to facilitate metabolic adaptation to hypoxia, allowing tumor cells to maintain energy production and survive in oxygen-deprived niches [19]. This hypoxic tolerance is frequently associated with resistance to treatment and aggressive tumor behavior [21].

#### Biological Interpretation and Implied Hypothesis

When expressed in terms of the above prior knowledge, the data-driven SDC4/NDUFA4L2 BMC model corresponds thus to the following hypothesis:

*In the clinical context, glioblastoma cells may sustain tumor proliferation and progression through two distinct, yet complementary, mechanisms: an invasive phenotype driven by SDC4-mediated signaling and hypoxic tolerance facilitated by NDUFA4L2. In tumors where either pathway is active, aggressive behavior is maintained. However, when both SDC4 and NDUFA4L2 are expressed at low levels, the tumor lacks both a robust invasive capability and an efficient hypoxic adaptation mechanism, resulting in reduced aggressiveness and improved long-term survival*.

In essence, this hypothesis, which we term *Double Defense Hypothesis*, posits that glioblastoma cells have two adaptive defense strategies: enhancing invasiveness or increasing hypoxic tolerance, to overcome hostile microenvironmental conditions. The absence of both strategies (i.e., low expression of SDC4 and low expression of NDUFA4L2) leaves tumor cells less equipped to proliferate and disseminate, thus correlating with better clinical outcomes.

Note that the Double Defense (DD) Hypothesis is significantly stronger than just the concatenation of existing prior knowledge:

- The DD hypothesis extends the notion that invasion and hypoxic adaptation capabilities are *individually important* for survival to the notion that, together, they are *determining factors* for survival.
- The DD hypothesis implies a specific “AND logic” of interaction between the two capabilities (long survival requiring the absence of both capabilities).
- The SDC4/NDUFA4L2 BMC implies specific transcriptomic threshold values for the interpretation of “low” expression in the DD hypothesis.

The data-driven BMC model of the interaction between the two genes and the survival phenotype has thus brought together two pieces of prior knowledge that, to our knowledge, had not been studied in a clinical context beforehand. Besides the DD hypothesis, the BMCs shown in Figure 5 suggests other new avenues for further investigation.

- Do cases that are misclassified by the SDC4/NDUFA4L2 classifier, in particular, the red ‘short survival’ dots on the lower left, present specific characteristics that could suggest why they were misclassified?
- Interestingly, the SDC4/NDUFA4L2 BMC is closely linked with another BMC, CHSY3/RASSF1 (Figure 5, upper right corner). CHSY3 is functionally related to SDC4, as CHSY3 is a key enzyme responsible for chondroitin sulfate biosynthesis, and its activity is thus directly relevant to the composition of the glucosaminoglycan chains that attach to SDC4 [22]. This raises several questions. First, as both SDC4 and CHSY3 are involved in cell motility, one might expect that the partner RASSF1 of CHSY3, just as NDUFA4L2, has a role in hypoxia adaptation. Indeed, its most well-studied isoform, RASSF1A, is known as one of the most frequently inactivated tumor suppressor genes [23], and has recently also been implicated in cellular adaptation to hypoxia [24]. Our data therefore suggests that this is also a role of RASSF1 in glioblastoma. Intriguingly, in our glioblastoma data, high overall RASSF1 expression is associated with *short*, instead of *long* survival, suggesting that the observed high RASSF1 expression comes from the non-A isoforms of RASSF1, whose significance remains to be explored.

#### Hypothesis Validation Using TCGA Data

As a simple first test of the Double Defense hypothesis, we apply it to the independent ovarian cancer dataset from TCGA [25]. Using the *ad hoc* choice of the 75th percentile of the SDC4/NDUFA4L2 expression in the TCGA cohort as the threshold between “low” and “high” expression, the Double Defense Hypothesis predicted 83 TCGA patients as long-term survivors and 71 and others as short-term survivors.

The resulting BMC in Figure 7a separates short and long survival in the TCGA much less accurately than the one in Figure 6. One of the reasons for poorer performance may lie in the differences between the two cohorts. For example, while the TCGA cohort contains “classic” primary glioblastomas, the Reifenberger et al. [12] cohort was specifically designed to capture extremes of the survival distribution. Furthermore, that cohort contains more recent, and more uniformly treated, set of patients, the majority receiving combined radiotherapy plus temozolomide (TMZ), where only a fraction in the older TCGA cohort were treated with TMZ.

**Figure 7:**
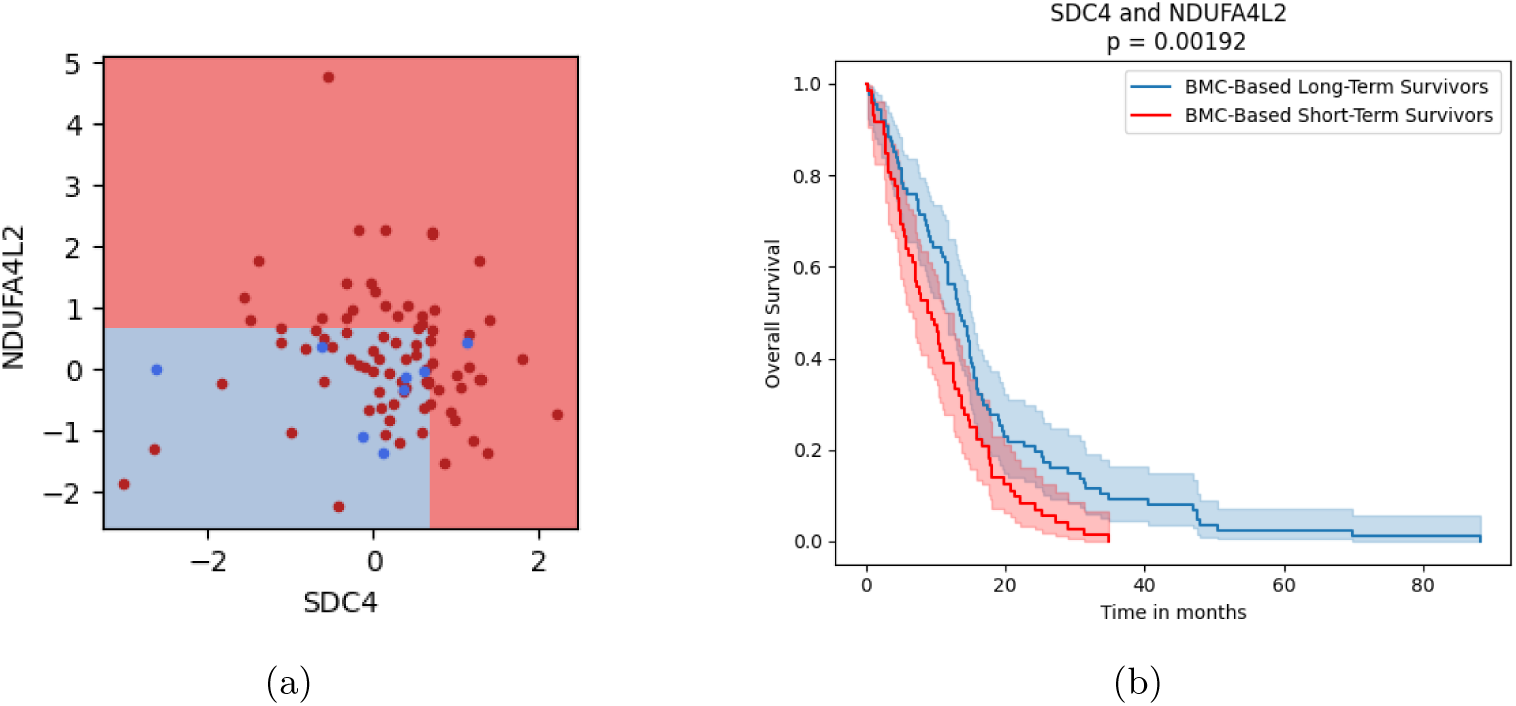
Application of the BMC to the independent TCGA glioblastoma cohort. (a) The BMC from the Reifenberger et al. [12] cohort, applied to the TCGA cohort. (b) Kaplan–Meier survival curves, stratified by BMC-predicted survival groups. The standard log-rank test indicates significantly longer survival in the predicted long-term survival group.

However, a statistical test using the actual survival data did confirm significantly longer survival of those patients that were predicted to survive longer (Figure 7b).

#### Options for Further Hypothesis Validation

The BMC structurally simple interaction model between two genes/processes allows several options for validation:

- **In Vitro Functional Studies:** Use glioblastoma cell lines to individually and jointly knock down SDC4 and NDUFA4L2 using siRNA or CRISPR-Cas9 approaches. Evaluate changes in cell migration, invasion (via Transwell or woundhealing assays), and proliferation under normoxic and hypoxic conditions. Assess whether simultaneous downregulation of both genes has a greater inhibitory effect than targeting either gene alone. Monitor downstream signaling pathways (e.g., PKC, RhoA, MAPK/ERK for SDC4, and hypoxia-responsive pathways for NDUFA4L2) using Western blotting and reporter assays.
- **In Vivo Xenograft Models:** Generate glioblastoma xenografts in immunocom-

promised mice with defined expression profiles of SDC4 and NDUFA4L2. Compare tumor growth, invasion, and overall survival between groups with low expression of both genes versus those with high expression of one or both.

- **Clinical Data Analysis:** Validate the association by correlating SDC4 and

NDUFA4L2 expression levels with clinical outcomes in independent glioblastoma cohorts and evaluate markers of hypoxia (e.g., HIF-1*α*) and invasion (e.g., MMP expression) in relation to the combined gene signature.

- **Multiomics Approaches:** Employ transcriptomic and proteomic profiling to de-

lineate the downstream networks modulated by SDC4 and NDUFA4L2. This may reveal synergistic interactions or compensatory mechanisms that contribute to tumor aggressiveness.

#### Therapeutic Implications

Agents targeting invasiveness and hypoxia adaptation in glioblastoma already exist. Cilengitide is a drug designed to disrupt SDC4-mediated adhesion and migration, yet failed in a Phase III trial to achieve the desired endpoint [26]. Apatinib effectively targets NDUFA4L2 [20] and was shown to have a certain level of antitumor activity [27].

If validated, the Double Defense Hypothesis suggests that effective glioblastoma therapies should simultaneously target both invasiveness and hypoxic adaptation to increase survival, in particular in those patients whose tumors exhibit simultaneous high expression of SDC4 and NDUFA4L2. Moreover, the joint low expression of these genes could serve as a prognostic biomarker to identify patients with inherently less aggressive tumors who might benefit from less intensive treatment regimens.

## 4 Formal Description of the Methods

This section is a formal description of the methods presented in this paper. It includes naïveBMC, whose detailed description has been omitted in Nikolayeva et al. [6] (Section 4.1), as well as the Preselection Algorithm and fastBMC (Section 4.2).

### 4.1 naïveBMC

This section breaks down the construction of BMCs (Section 4.1.1) and the various stages that make up the naïveBMC (Section 4.1.2 and Section 4.1.3).

#### 4.1.1 Classification Setting and Bivariate Monotonic Classifiers

Formally, the input data in [6] is given as a vector of values for *m* ∈ **N**≥_1_ transcriptomic features, for each of *n* ∈ **N**≥_1_ patients. Each patient also has a disease label in {0, 1}. The problem aims to predict, for a new patient, the disease label from the transcript values.

Nikolayeva et al. [6] used *bivariate monotonic (binary) classifiers (BMCs)*, i.e., functions *f* : ℝ^2^ → {0, 1} that are monotonic in both dimensions.^1^ Any function that increases monotonically in *x* or −*x*, and in *y* or −*y* is a BMC. An *optimal* BMC *f* is one such that *f* minimizes the *classification error* (ER_full_) of the input with respect to its labels 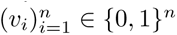, that is, 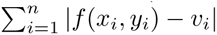.

Since a ER_full_-minimal *f* might not be unique, we follow the approach of Nikolayeva et al. [6] and define an *optimal* BMC as one that classifies the most points in **R**^2^ as 1. Formally, a ER_full_-minimal BMC *f* ^∗^ constructed over *S* ⊂ **R**^2^ is considered optimal if and only if for all ER_full_-minimal BMCs *f* constructed over *S* holds that |{*x* ∈ **R**^2^ | *f* ^∗^(*x*) = 1}| ≥ |{*x* ∈ **R**^2^ | *f* (*x*) = 1}|.

The CE of the BMC in Figure 1 is 1, as exactly one blue triangle is within the red area and no red point falls in the blue area. The red area cannot be further increased without increasing the ER_full_. Therefore, this BMC is optimal.

A ER_full_-minimal BMC for *n* points can be efficiently computed by dynamic programming in time Θ(*n* log^2^ *n*) [7]. Since this algorithm only constructs monotonically *increasing* BMCs, we compute four different ER_full_-minimal BMCs for each transcriptomic feature pair – one for each possible orientation of each of the two axes. We then choose a ER_full_-minimal BMC among these four options.

Note that although BMCs can be *described* as decision trees, common greedy algorithms for decision trees typically do not *identify* classifiers such as those in Figure 1. This may explain the poorer performance of the decision trees that were computed in Nikolayeva et al. [6] for comparison.

#### 4.1.2 Ensemble Bivariate Monotonic Classifiers

Nikolayeva et al. [6] explain how to combine multiple BMCs into an ensemble classifier of at most *k*_max_ BMCs. Such an *ensemble BMC* is a set of *k* ∈ **N**≥_1_ BMCs, whose predicted label is the majority of the binary labels predicted by the *k* BMCs.^2^

The classification performance of any given BMC is estimated by removing a patient in turn from the input, training a BMC on the reduced dataset, and evaluating the absolute difference (0 or 1) between the predicted and the true label of the removed patient (this is called *leave-one-out cross-validation* (LOOCV)). The sum of these values is the BMC’s *LOOCV error*, or *ER_loocv_*(Section 4.1.3).

The approach of Nikolayeva et al. [6] is divided into three phases.

- **Phase 1: Estimate the Optimal Number *k*^∗^ of BMCs** For each *k* ∈ [1*, k*_max_] ∩ **N**, estimate the classification performance of an optimal ensemble BMC, as follows. For each patient in turn, an ensemble BMC is constructed using all patients except the one evaluated (using a LOOCV). The error rates of these BMCs are calculated, and the top BMCs are selected for various values of *k*. The value of *k* whose model has the lowest ER_loocv_ is chosen as the optimal number *k*^∗^ of BMCs.
- **Phase 2: Estimate Classification Performance with *k*^∗^ BMCs** The second phase evaluates the performance of the ensemble BMC constructed with the *k*^∗^ BMCs determined in Phase 1. Patient predictions are made using an ensemble BMC, and misclassification rates, calculated with LOOCV, are recorded. The performance of the ensemble BMC is quantified by estimating its error rate, calculated as the proportion of patients misclassified.
- **Phase 3: Constructing the Ensemble Classifier** To create the final ensemble BMC, we re-evaluate all BMCs using a LOOCV across all patients. This ensures robustness. The top *k*^∗^ performing BMCs are then chosen to build the ensemble. This ensemble BMC aims to achieve the highest possible AUC score^3^ by incorporating the most informative BMCs.

#### 4.1.3 Prediction Error Estimate by Leave-One-Out Cross-Validation

Nikolayeva et al. [6] used LOOCV to estimate the prediction error of a BMC from a dataset 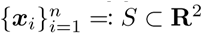 of gene expression measurements for *n* patients, with labels 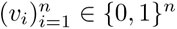. For a given pair of genes, for each *i* ∈ [1*, n*] ∩ **N**, they construct the optimal BMC *f* for *S* \ {***x****_i_*} and compute its prediction error |*f* (***x****_i_*) − *v_i_*| =: *ε_i_*. They use the *LOOCV error* of the optimal BMC for *S* as 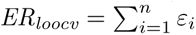 to estimate the predictive performance of the BMC on new data. Evaluating the ER_loocv_ of each BMC is computationally expensive, as it requires the construction of a new BMC for each left out patient. By computing the ER_loocv_ of all *m* pairs of transcriptomic features, the overall cost is *mn*, since, for each pair of transcriptomic features, each of the *n* patients needs to be left out in turn. In contrast, when using ER_full_, the expensive LOOCV only needs to be performed for a subset of *ℓ* ≤ *m* pairs of transcriptomic features. This results in an overall cost of *ℓn* + *m*, where the term *m* accounts for the evaluation of the ER_full_. If *ℓ* ≪ *m*, then the overall cost of this new approach is much lower than that of the naïve approach.

### 4.2 The Preselection Algorithm and fastBMC

This section gives a detailed presentation of the additions to naïveBMC. These include the proof a property (Section 4.3), enabling the identification and selection of the bestperforming BMCs (Section 4.3.1). And it ends with the modification of naïveBMC, into its new version fastBMC (Section 4.3.3).

### 4.3 The Classification Error is a Lower Bound for the LOOCV Error

We prove in Theorem 1 that ER_full_ is a lower bound for ER_loocv_. Theorem 1 shows that any point ***x*** ∈ **R**^2^ that is correctly predicted by an optimal BMC constructed on a set of *n* − 1 points is also correctly predicted by an optimal BMC constructed over the same *n* − 1 points *and* ***x***.

In order to state the theorem and its proof concisely, we introduce some notation. Let *S* ⊂ **R**^2^ be a set of data points. For all *U* ⊆ *S*, let C(*U* ) denote the set of all optimal BMCs over *U* . Furthermore, for all *U, V* ⊆ *S* and all BMCs *C* ∈ C(*U* ), let MC(*C, V* ) denote the number of misclassified points among *V* according to *C*, that is, its ER_full_. **Theorem 1.** *Let S* ⊂ **R**^2^*, and let* ***x*** ∈ *S. Furthermore, using the notation above, let C_S_* ∈ C(*S*) *and C_S_*_\{_***_x_***_}_ ∈ C(*S* \ {***x***})*, and assume that* ***x*** ∈ *S is correctly classified by C_S_*_\{_***_x_***_}_*. Then* MC(*C_S_, S*) = MC(*C_S_, S* \ {***x***}).

*Proof.* We show the equation to be proved via the following two cases.

**Case MC(*C_S_, S*) *≥* MC(*C_S_, S \ {x}*).**

This inequality holds because eliminating a data point evaluated by *C_S_* only keeps or reduces the number of misclassified points.

**Case MC(*C_S_, S*) *≤* MC(*C_S_, S \ {x}*).**

Since *C_S_*_\{_***_x_***_}_ is optimal (with respect to ER_full_) to classify *S* \ {***x***}, we have MC(*C_S_, S* \ {***x***}) ≥ MC(*C_S_*_\{_***_x_***_}_*, S* \ {***x***}). Since ***x*** is correctly classified by *C_S_*_\{_***_x_***_}_ by assumption, we have MC(*C_S_*_\{_***_x_***_}_*, S* \ {***x***}) ≥ MC(*C_S_*_\{_***_x_***_}_*, S*). Moreover, as *C_S_*is optimal for classifying *S*, it holds that MC(*C_S_*_\{_***_x_***_}_*, S*) ≥ MC(*C_S_, S*). By transitivity, this case follows.

**Conclusion.** Combining both cases concludes the proof.

#### 4.3.1 The Preselection Algorithm

Building on Theorem 1, we propose a Preselection Algorithm (Algorithm 1) that identifies a set of top-performing pairs of transcriptomic features for the BMCs (Section 2.1). We explain the inner workings of the algorithm, that is, how the ER_full_ threshold of the Preselection Algorithm changes over time and how this affects how many BMCs are eliminated.

#### 4.3.2 Formal Implementation

Given an integer *k*, the Preselection Algorithm returns a set of BMCs that contain at least *k* pairwise disjoint pairs of transcriptomic features *(feature pairs)*. In order to reduce the expensive LOOCV computation, the main idea is to determine and maintain a ER_full_ threshold *t*, which allows removing BMCs based on their ER_full_ instead of their ER_loocv_. On a high level, the Preselection Algorithm operates in three phases:

1. Evaluate the ER_full_ of all BMCs for all feature pairs and sort them with respect to increasing ER_full_ (lines 1 to 6).
2. Evaluate the ER_loocv_ of the BMCs with the lowest ER_full_ until the output contains at least *k* disjoint feature pairs, while also determining a threshold ER_full_ (lines 7 to 22).
3. Evaluate the ER_loocv_ of the remaining BMCs, adding better ones to the output, and update *t* (lines 23 to 41).

For Phase 1, we note that ties in the ER_full_ of different classifiers are broken arbitrarily (e.g., using lexicographic or random ordering).

Phase 2 chooses the ER_full_ threshold *t* as the maximum ER_loocv_ among all BMCs evaluated, which are stored in a search tree *T* . In this phase, we also store the number of disjoint feature pairs among the evaluated BMCs in the variable *d*. With each new BMC *C* that is evaluated, we iterate over all BMCs in *T* and determine their feature pairs. If the features of *C* do not appear in any existing feature pair, we increase *d* by 1, as we found a new feature pair. Otherwise, we proceed with the next iteration. This naïve method of checking the number of pairs of disjoint features could easily be improved. However, since it corresponds to the method of Nikolayeva et al. [6] to select the best feature pairs, it is kept for consistency reasons.

In Phase 3, the remaining BMCs are considered in increasing order according to their ER_full_. Due to Theorem 1, if a BMC has a ER_full_ greater than the current threshold *t*, its ER_loocv_ is also at least as bad. As the BMCs are ordered by their ER_full_, we immediately skip the evaluation of any additional BMCs. Otherwise, we evaluate the ER_loocv_ of the BMC. During this evaluation, since we have a threshold *t*, if we see that the ER_loocv_ is worse than *t*, we stop the LOOCV and ignore the BMC. Otherwise, we add it to *T* . As *T* is increased, it might be the case that we can remove the BMCs that are too bad by now, recalling that we keep or eliminate all BMCs of the same ER_loocv_. To this end, we check whether if we remove all BMCs with the worst ER_loocv_ (which is *t*), we still have at least *k* disjoint feature pairs. We check this in the same way as in Phase 2. If this is the case, we remove all BMCs with a ER_loocv_ of *t*, and we update *t* to the new worst ER_loocv_ among all BMCs that we still keep.

At the end of the Preselection Algorithm, we return all BMCs that were kept. Note that after a run of the Preselection Algorithm, the BMCs are partitioned into three types: (1) those that are eliminated without computing their ER_loocv_, (2) those for which the ER_loocv_ is (if possible, partially) computed but *not* selected in the output, and (3) those for which the ER_loocv_ is computed and that *are* part of the output. For example, for the dengue dataset (Figure 8), at the end of the algorithm, more than 97 % of all BMCs have been eliminated without computing their ER_loocv_, while only a minimal percentage of BMCs are kept among the ones that are evaluated.

**Figure 8:**
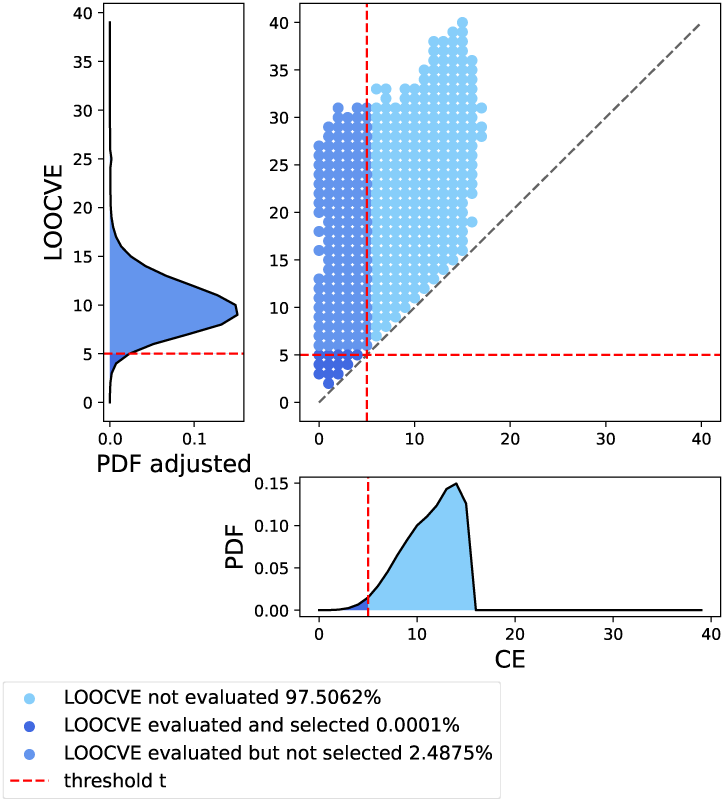
The reduction in the number of the ER_loocv_ evaluations using the Preselection Algorithm (Algorithm 1) on the dengue dataset (Section 3.1.1). Each point represents a set of potentially many BMCs with the ER_full_ and ER_loocv_ corresponding to the coordinates. The graphs along both axes represent the densities (probability density functions; PDFs) of the BMCs. The PDF at the bottom ranges over the whole dataset, the PDF at the left only over the BMCs that are evaluated (approximately 5 % of the overall data). The vertical dashed line represents the cutoff point at which no further ER_loocv_ evaluations were necessary. The horizontal dashed line shows the cutoff point beyond which no additional BMCs were selected among those whose ER_loocv_ was computed. Note that the update of the ER_full_ threshold *t* is not shown.

#### 4.3.3 fastBMC

The fastBMC algorithm is an extension of naïveBMC that incorporates the Preselection Algorithm. The Preselection Algorithm is called within each of the three phases of naïveBMC mentioned in Section 4.1.2. Each of these phases conducts an exhaustive LOOCV. During each of these LOOCVs, immediately after every sample that is excluded for testing, the Preselection Algorithm is called. This means that the Preselection Algorithm is called multiple times per phase. The reason for calling the Preselection Algorithm repeatedly is that this ensures that we do not bias the overall LOOCV with respect to which patient we leave out. In more detail, the Preselection

##### Algorithm 1

The Preselection Algorithm for identifying optimal BMCs. Let *m* ∈ **N**≥_1_ be the number of feature pairs, *n* ∈ **N**≥_1_ the number of patients, and let 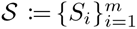 be the feature pairs with their respective labels 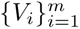 such that, for all *i* ∈ [1*, m*] ∩ **N**, it holds that *S_i_* ∈ (**R**^2^)*^n^* and *V_i_* ∈ {0, 1}*^n^*. Further, let *k* ∈ **N**≥_1_ with *k* ≤ *m* be given. The Preselection Algorithm returns a set of BMCs with the lowest ER_loocv_ among S containing at least *k* disjoint feature pairs, if possible.

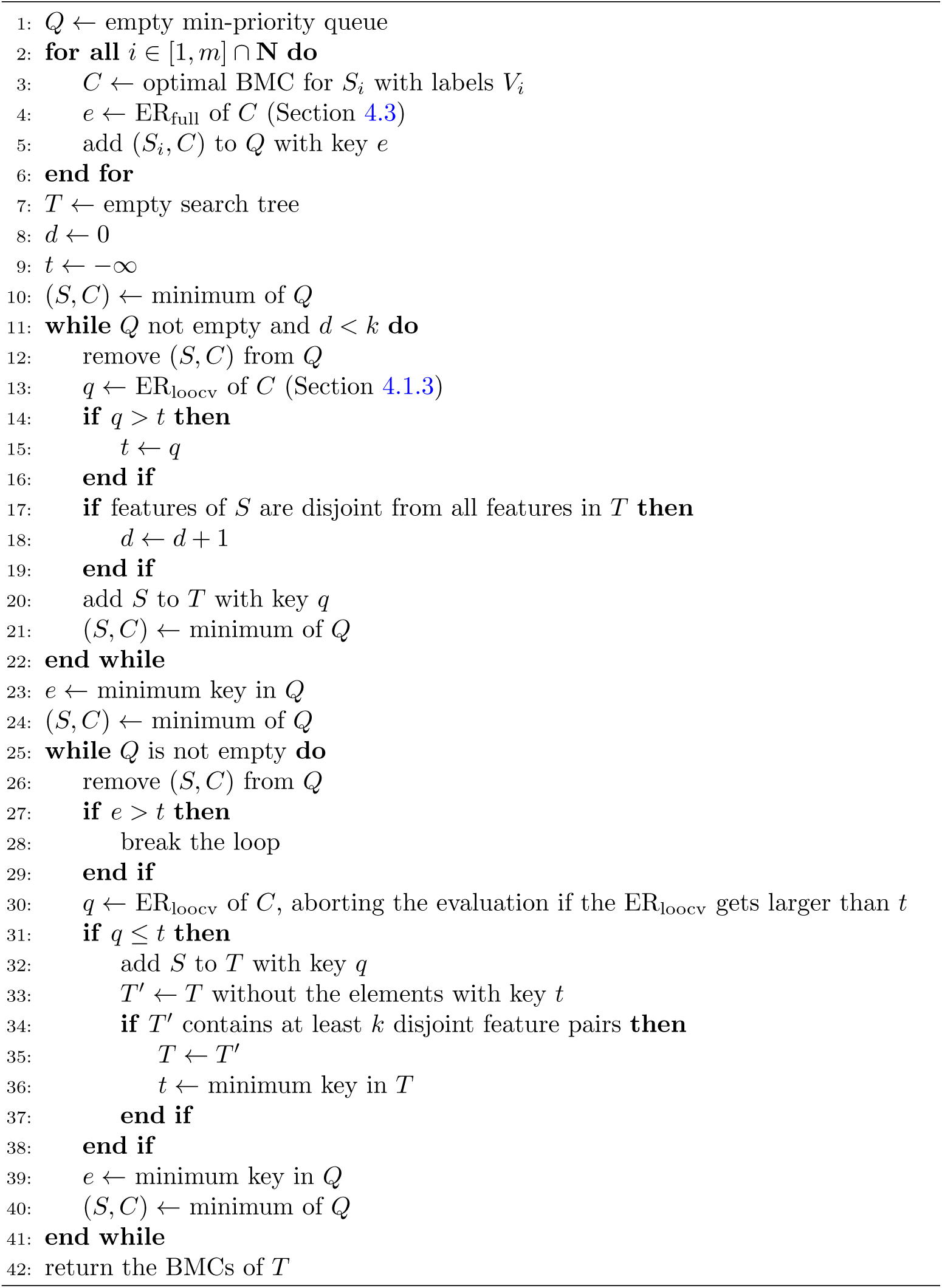

Algorithm does not introduce bias into the subsequent computations because it does not have access to the data point that is used for testing.

## 5 Discussion

Our empirical evaluation on three clinical datasets suggests that fastBMC produces models that perform similarly to naïveBMC, in drastically less time, thus allowing the analysis of significantly larger sets of features, which, in turn, is sometimes critical to discover better classifiers.

In this article, we introduce an extension of an existing method by Nikolayeva et al. [6], which we call fastBMC. This extension is based on a Preselection Algorithm (Algorithm 1) that operates on a set of optimal bivariate monotonic classifiers *(BMCs)* for transcriptomic feature pairs and efficiently determines those with the lowest leave-one-out cross-validation error. Such sets are invaluable for constructing optimal ensembles of BMCs, which are not only highly accurate, but also easily interpretable. When combining the Preselection Algorithm with the approach by Nikolayeva et al. [6] to compute BMCs, we empirically find that for datasets of 250 and more features, fastBMC is at least 15 times faster than the original approach (naïveBMC), and that this speed-up is also required in our empirical evaluation to discover optimal classifiers in reasonable computation time, in two out of three datasets. Our study acknowledges that an increasing number of features does not necessarily translate to better classification performance, and in fact, when the number of features far exceeds the number of samples, the curse of dimensionality can lead to overfitting, making it easier to fit a model to noise rather than meaningful patterns. This concern is particularly relevant when using fastBMC, which can handle larger numbers of features and thus increase the risk of overfitting. Compared with classical classification methods, fastBMC performs as well as the other methods. The benefit lies in the interpretability and visualization of the model. Thus, fastBMC represents a considerable advance for the discovery of BMCs especially in transcriptomic but potentially also data-richer biological applications, where even more features are available. The software we used for this study is the first open source library, in Python, that allows the discovery of BMCs and their ensembles. It is available at https://github.com/oceanefrqt/fastBMC. Direct interpretation of the identified glioblastoma BMC SDC4/NDUFA4L2 in the context of prior knowledge immediately suggested a novel plausible and experimentally testable clinical hypothesis and downstream implications for glioblastoma treatment (Section 3.4).

It is important to temper our observation with an acknowledgment of the dynamic complexity inherent in glioblastoma. These tumors are characterized by extensive intratumoral heterogeneity and rapid clonal evolution, factors that may not be fully captured by a single pretreatment transcriptomic snapshot. In practice, the aggressive and adaptive nature of glioblastoma means that small subpopulations of cells with distinct gene expression profiles could emerge over time, potentially altering the tumor behavior despite a favorable initial baseline signature. Therefore, even if the SDC4/NDUFA4L2 signature provides valuable prognostic information, it should be viewed as one piece of a larger and more complex puzzle, in which the two physiological aspects brought together by the BMC may also, biologically, be more closely related than initially thought [28].

Perhaps due to these complexities, our initial validation in the TCGA cohort could confirm a statistical association of SDC4/NDUFA4L2 with survival, but not the originally much stronger association in a more recent and more homogeneous cohort. Future studies in other cohorts employing single cell sequencing and longitudinal monitoring may help characterize those subtypes of glioblastoma in which the association can be observed, how stable the SDC4/NDUFA4L2 signature remains over time, and how they integrate with other dynamic molecular changes that drive the progression of glioblastoma.

Motivated by the performance and interpretability in our glioblastoma case study, our methodological efforts are now directed towards further refinement of fastBMC and exploration of its applicability to biological and non-biological datasets. Currently, BMCs produce binary output. To enhance the practical value of BMCs, it would be interesting to estimate the probability of correctness for each prediction. This probability could be defined, for example, on the basis of the distance of a point from the classification boundary by a method such as Platt scaling [29]. Platt scaling is a calibration technique used in machine learning to convert the output of a classifier into well-calibrated probability scores, typically by applying a logistic function to model predictions.

Alternatively, a monotonic multiclass model could be used. Using any inherent order of training data, a multiclass approach represents a form of ordinal classification, typically used in bioinformatics for tasks such as cancer staging or prediction of disease progression. Beyond biology, ordinal classification is useful in other domains such as satisfaction surveys [30], bank failure prediction [31], and quality control [32].

## Declarations

### Ethics approval and consent to participate

Not applicable

### Consent for publication

Not applicable

### Availability of data and materials

The leukemia and glioblastoma datasets are both publicly available, as mentioned in the Section 3.1.1. The dengue dataset is available per request to the authors. The TCGA validation data are publicly available through cBioPortal (https://www.cbioportal.org/study/summary?id=gbm tcga pan can atlas 2018).

The code to compute BMCs is available at https://github.com/oceanefrqt/ fastBMC.

### Authors’ Contributions

OF designed and implemented the computational code and performed data analysis. MK, CD, and OF developed the theoretical framework and proof of the preselection algorithm. BS investigated the biological hypotheses resulting from computational analysis. OF and MK wrote the manuscript, with significant input from the other authors. CD and BS oversaw the project, provided critical feedback, and refined the manuscript. All authors reviewed and approved the final manuscript.

## Acknowledgements

We thank the patients and their families, without whom this work would have not been possible.

## Footnotes

1 Given a set of points 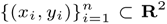, a *monotonically increasing* classifier (in *x* and in *y*) is any function *f* : **R**^2^ → {0, 1} such that for all *i, j* ∈ [1*, n*] ∩ **N** it holds that if *xi* ≤ *xj* and *yi* ≤ *yj* , then *f* (*xi, yi*) ≤ *f* (*xj, yj* ).

2 In the case of a tie, the label is chosen to be 1.

3 AUC stands for Area Under the Curve. The AUC ROC metric, abbreviated as AUC, consists of calculating the area under the Receiver Operating Characteristic curve, which plots all the values of the pair (1-Specificity, Sensitivity) according to the classification threshold.

